# Customized de novo mutation detection for any variant calling pipeline: SynthDNM

**DOI:** 10.1101/2021.02.10.427198

**Authors:** Aojie Lian, James Guevara, Kun Xia, Jonathan Sebat

## Abstract

**Motivation:** As sequencing technologies and analysis pipelines evolve, DNM calling tools must be adapted. Therefore, a flexible approach is needed that can accurately identify de novo mutations from genome or exome sequences from a variety of datasets and variant calling pipelines.

**Results:** Here, we describe SynthDNM, a random-forest based classifier that can be readily adapted to new sequencing or variant-calling pipelines by applying a flexible approach to constructing simulated training examples from real data. The optimized SynthDNM classifiers predict de novo SNPs and indels with robust accuracy across multiple methods of variant calling.

**Availability:** SynthDNM is freely available on Github (https://github.com/james-guevara/synthdnm)

**Supplementary information:** Supplementary data are available at *Bioinformatics* online.

## 1. Introduction

Genome wide analysis of *de novo* mutation by whole genome sequencing (WGS) is being applied in a variety of settings, from small studies of individual families(Kong et al., 2012) to cohorts of several thousand genomes(An et al., 2018; Zhou et al., 2019). Thus, there is a need for variant classification methods that can detect DNMs in a variety of datasets with high accuracy. When new sequencing protocols, datasets or data processing pipelines are introduced, the process of assembling new training data is labor intensive.

## 2. Methods

We have developed SynthDNM, a software package written in python that enables optimal DNM detection in any new dataset or variant calling pipeline by rapid optimization of the classifier using real data. SynthDNM requires a Fam and VCF file as input.

## 3. Classifier

The default SynthDNM classifier is designed for GATK variant calls on Illumina sequence data. In addition SynthDNM can be re-trained for any new dataset or data type by re-implementing the training procedure with a new VCF file contain jointly-called variants on trio families. SynthDNM creates a large set of simulated DNMs from real variants by swapping parents and offspring between families (Fig. 1A). When private inherited variants in offspring are matched to the surrogate (unrelated) parents that have homozygous reference genotypes, they are classified as “synthetic DNMs”. This ensures that simulated DNMs have the same properties as other ultra-rare germline variants. Lastly, since the majority of putative DNMs in the original trio represent errors (either a false-positive variant in the offspring or a false-negative genotype in a parent), all putative de novos are assigned to the negative training set. The default classifier of SynthDNM, was trained on a jointly-called VCF from >30X Illumina sequence from the Simplex Collection (SSC) WGS dataset (SSC-GATK classifier).

**Fig 1.**
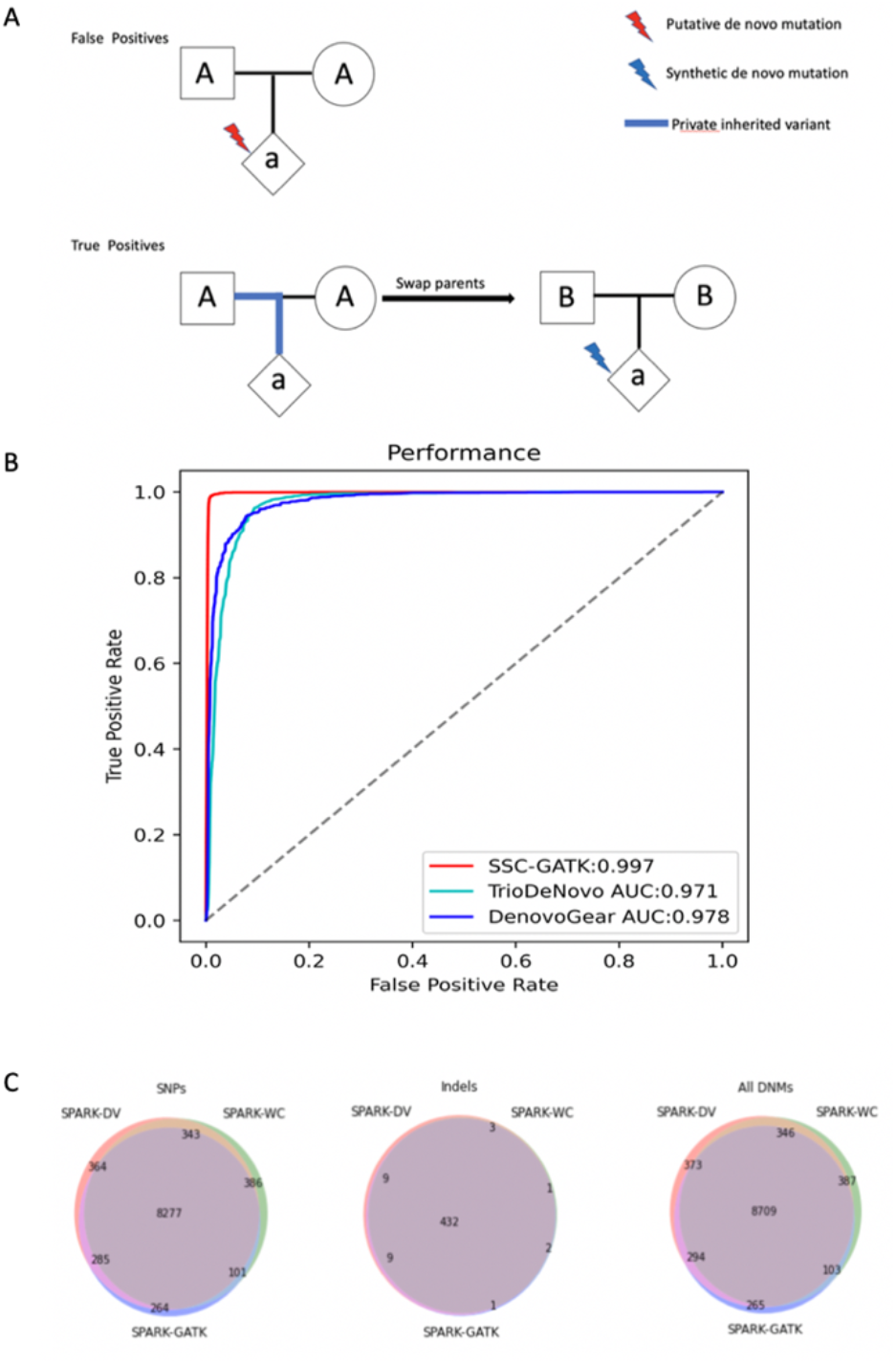
(A)the methods of generating “synthetic DNMs”. (B) ROC curves of SSC-GATK classifier, Triodenovo and DenovoGear. All classifiers were predicted on the same 3787 individuals. (C) Different DNM prediction performance of the different calling methods.

## 4. Performance results

SSC-GATK DNM calls intersecting with exons were validated using a set of de novo mutations detected by Whole Exome Sequencing (WES) of the same samples (Iossifov *et al*., 2014). We then compared the accuracy of SSC-GATK to DNM calls made with DeNovoGear and TrioDeNovo on the same SSC data. The accuracy of SSC-GATK classifier was improved (AUC = 0.997) compared to DeNovoGear (AUC = 0.978) and TrioDeNovo (AUC = 0.971) (Fig. 1B).

In addition, we compared the quality of SSC-GATK calls to DNMs reported on the SSC WGS dataset in two previous studies(Werling *et al*., 2018; Turner *et al*., 2017). In this case, “false positive” DNMs were defined as exonic DNM calls in the WGS dataset that were not detected in a previous exome sequencing study. This definition of FDR is an overestimate, since a subset of non-validated DNMs may be true positives, but this metric allows us to assess accuracy of the calls sets relative to each other. The TPR and F1 of SSC-GATK classifier(0.97 and 0.82) is higher than for the other two call sets (Werling:0.83,0.77;Turner:0.30,0.40),which suggests that a single optimized method can provide greater sensitivity and overall accuracy compared to the intersection of multiple methods or set of stringent filtering heuristics.

In recent years, a number of new variant-calling platforms have been introduced(Poplin *et al*., 2018). Each uses a unique set of algorithms for evaluating variant and genotype quality, and for these new platforms our classifier is not compatible. We therefore evaluated whether SynthDNM could make a consistent set of DNM calls across multiple variant calling platforms applied to a single dataset, SPARK exome sequencing study(Feliciano *et al*., 2019), including GATK(SPARK-GATK), Deep_Variant(SPARK-DV), and We Call(SPARK-WC). Across a variety of variant calling platforms, the customized synthDNM classifiers, for the three platforms made consistent DNM predictions (Fig. 1C). For SNPs, 82.6% (N=8277) of variants overlapped across all three different platforms; For indels, 94.5% (N=432) of variants overlapped across all three different platforms. To further confirm the accuracy of our synthDNM calls, we compared the de novo SNP and indel calls on the SPARK dataset to a set of validated de novo mutations that were confirmed by Sanger sequencing in the same cohort in a previous pilot study (Feliciano et al., 2019) to test our classifier. For SNPs, Spark-GATK could recall 112/114(98.2%) de novo snps; Spark-DV could recall 109/117(93.2%) de novo snps; Spark-WC could recall 100/108(92.6%) de novo snps. For Indels, Spark-GATK could recall 106/107(99.1%) de novo indels; Spark-DV could recall 84/84(100%) de novo indels; Spark-WC could recall 68/69(98.6%) de novo indels.

## 5. Conclusion

Optimized SynthDNM classifiers detect de novo SNVs and indels with high accuracy across multiple dataset and variant calling pipelines.

## Acknowledgements

We thank the members of the Centre for Medical Genetics, Central South University and Department of Psychiatry, University of California San Diego for their valuable discussion regarding this work.

## References

An, J.-Y. et al. (2018) Genome-wide de novo risk score implicates promoter variation in autism spectrum disorder. Science, 362.

Feliciano, P. et al. (2019) Exome sequencing of 457 autism families recruited online provides evidence for autism risk genes. NPJ Genomic Med., 4, 19.

Iossifov, I. et al. (2014) The contribution of de novo coding mutations to autism spectrum disorder. Nature, 515, 216–221.

Kong, A. et al. (2012) Rate of de novo mutations and the importance of father’s age to disease risk. Nature, 488, 471–475.

Poplin, R. et al. (2018) A universal SNP and small-indel variant caller using deep neural networks. Nat. Biotechnol., 36, 983–987.

Turner, T.N. et al. (2017) Genomic patterns of de novo mutation in simplex autism. Cell, 171, 710–722.e12.

Werling, D.M. et al. (2018) An analytical framework for whole genome sequence association studies and its implications for autism spectrum disorder. Nat. Genet., 50, 727–736.

Zhou, J. et al. (2019) Whole-genome deep-learning analysis identifies contribution of noncoding mutations to autism risk. Nat. Genet., 51, 973–980.

