## SupplementaryMaterials for "Customized de novo mutation detection for any variant calling pipeline: SynthDNM"

*Bioinformatics*, YYYY, 0–0
doi: 10.1093/bioinformatics/xxxxx
Advance Access Publication Date: DD Month YYYY
Applications Note

*Supplementary Material*

### Customized de novo mutation detection for any variant calling pipeline: SynthDNM

**Aojie Lian[1,2], James Guevara[2], Kun Xia[1], &Jonathan Sebat[2,*]**

1.Center for Medical Genetics & Hunan Key Laboratory of Medical Genetics, School of Life Sciences, Central South University, Changsha, Hunan, China.
2. Department of Psychiatry, University of California San Diego, La Jolla, California, United States of America.

Associate Editor: XXXXXXX

Received on XXXXX; revised on XXXXX; accepted on XXXXX

#### Introduction

Varioius informatics tools have been developed to detect DNMs. VarScan(Koboldt et al., 2009) is a mutation caller for targeted, exome and whole-genome resequencing data on different platform like illumina, SOLiD, Roche/454, and similar instruments. VarScan2(Koboldt et al., 2012), which was subsequently updated, can detect somatic mutations and copy number variations in cancer exome sequencing. Polymutt(Li et al., 2012) is a program implemented a likelihood-based framework for calling variants and detecting de novo point mutation in families. This method is based on the Elston-Stewart peeling algorithm to evaluate specific genetic variants, individual genotypes, and DNMs. Later, an improved version of Polymutt, called Triodenovo(Wei et al., 2015) based on the Bayesian model was introduced. This method separates the prior mutation rate from evaluation of the likelihood of the data and applies simulation and real data to prove that this method has higher sensitivity and specificity than other methods. DeNovoGear (Ramu et al., 2013) is a tool to detect somatic and germline de novo mutations on next-generation sequencing data, which use likelihood-based error modeling to reduce the false positive rate. novoCaller (Mohanty et al., 2019) is a Bayesian variant calling algorithm from pedigree and population sequence data. We previously published forestDNM (Michaelson et al., 2012) a random forest algorithm implemented in R to detect de novo single nucleotide mutations from family WGS data.

The methods described above are dependent on the quality and size of available training data, but the currently available training sets consist of a few thousand validated and invalidated DNMs derived from existing Illumina datasets and called with a pipeline based on GATK best-practices(Gardner et al., 2019; Werling et al., 2018; Brandler et al., 2018).When new sequencing protocols, datasets or data processing pipelines are introduced, the process of assembling new validated training data is lengthy. SynthDNM is a software package for optimizing DNM filtering for a variety of datasets and variant calling pipelines using simulated DNMs creating from variant calls in real data.

#### Methods

*Software Availability*

*SynthDNM* source code and documentation is hosted on GitHub

- Source Package: https://github.com/james-guevara/synthdnm
- Full Documentation: https://github.com/james-guevara/synthdnm/wiki

*SynthDNM Workflow*

SynthDNM is an open-source application written in Python that requires a pedigree (.fam or .ped) file(Purcell *et al.*, 2007) and a variant call format (VCF) file as input. There are two options to predict DNMs via SynthDNM.

1. To use the original classifiers that we provide on GitHub, the input VCFs must be called by GATK and recalibrated using VQSR. We provide four classifiers: an SNV autosomal classifier, an indel autosomal classifier, and an SNV classifier for the male sex chromosomes, and an indel classifier for the male sex chromosomes.

First, the software performs a preprocessing step to extract all the putative de novo mutations and their corresponding features from the VCF. Then, the classifiers predict DNMs using those features and output the results to a Browser Extensible Data (BED) file.

After obtaining all the putative de novo mutations, one should/may filter variants located in the regions that contain short tandem repeats (STRs) and segmental duplications, as well as variants located in unmapped regions, by using a BED intersection tool(Quinlan and Hall, 2010). Then one should filter out the common variants. In our case, we use ANNOVAR to obtain allele frequencies from the 1000 Genomes and gnomAD databases and filter out variants with frequencies greater than 1%. In our case, we also filter out predicted DNMs that appear in more than one individual (recurrent DNMs). Finally, we remove any individuals whose DNM count is an outlier.

**Fig.S1**

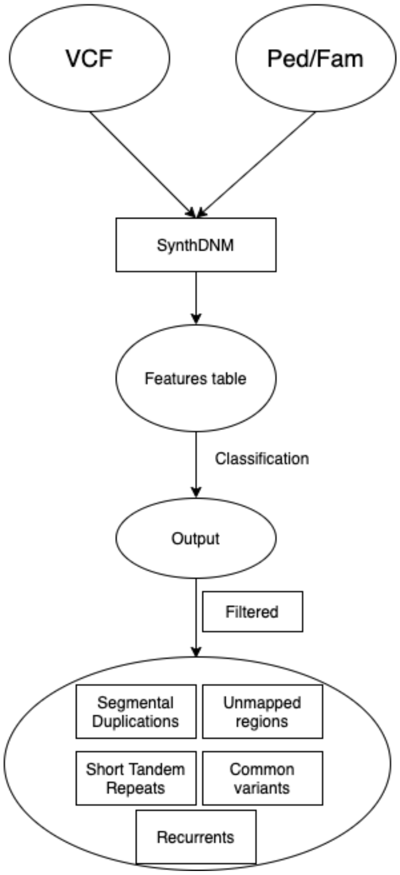


Fig.S1 SynthDNM workflow. SynthDNM requires a VCF and a fam file as input. The first step is to obtain putative DNMs from the VCF file. Then extract features from the VCF files pertaining to fields obtained from GATK VQSR for each sample. Finally, SynthDNM will classify all the variants after filtering procedure to remove variants located in regions that contain short tandem repeats and segmental duplications, and unmapped regions, and filter out common variants (variants with allele frequency > 1% in 1000 Genomes or gnomAD).

2.GATK is the industry standard for calling variants in germline DNA data, but other variant-calling software, such as DeepVariant or WeCall, which may result in features that the aforementioned classifiers cannot use for prediction. It may also be the case that due to changes in the GATK variant-calling pipeline (or in the DNA sequencing), our classifiers may perform below par. Therefore, we provide a method for creating classifiers using one's own dataset of choice (provided that the dataset contains parent-offspring trios). This method is essentially equivalent to the method we use to create our own classifiers, and the method is as follows:

a) Obtain *all* putative DNMs from the offspring in the dataset. In our case, using a WGS dataset as the input, this number was approximately 2800 for SNVs and approximately 370 for indels, per individual. This set of DNMs will be used as the negative training examples for the classifiers. (While these will also contain the real DNMs, these will be a miniscule fraction of all putative DNMs, as the numbers of real SNV and indel DNMs are around 60 and 10 on average per individual.)

b) To obtain the positive training examples, we first obtain private, inherited variants from the cohort of interest. As a preprocessing step, we use BCFtools to filter out multiallelic variants and to annotate the VCFs with an ID indicating relevant variant information (chromosome, position, reference allele, and alternate allele). We then use PLINK to obtain allele count information for each variant for inherited variants. Using this information, we then create a VCF with only the private, inherited variants. These private, inherited variants will be used as the "synthetic DNMs" that make up the positive training dataset.

c) The remaining steps are to first create a .fam/.ped file where all the parents have been swapped, and then use this swapped .fam/.ped file and the VCF with the private, inherited variants as inputs to the synthDNM extraction method. Since a private, inherited variant will look like a de novo variant when the .fam/.ped file is swapped, this process will produce putative DNMs, which will form the positive training examples that we feed into the classifiers.

d) Now that we have the entire training dataset, we can train our classifier. We use the random forest classifiers from the scikit-learn library for training and run a grid search and cross-validation step, varying a number of hyperparameters discussed in the next section. Once these steps are completed, a new set of classifiers will have been created.

**Fig.S2**

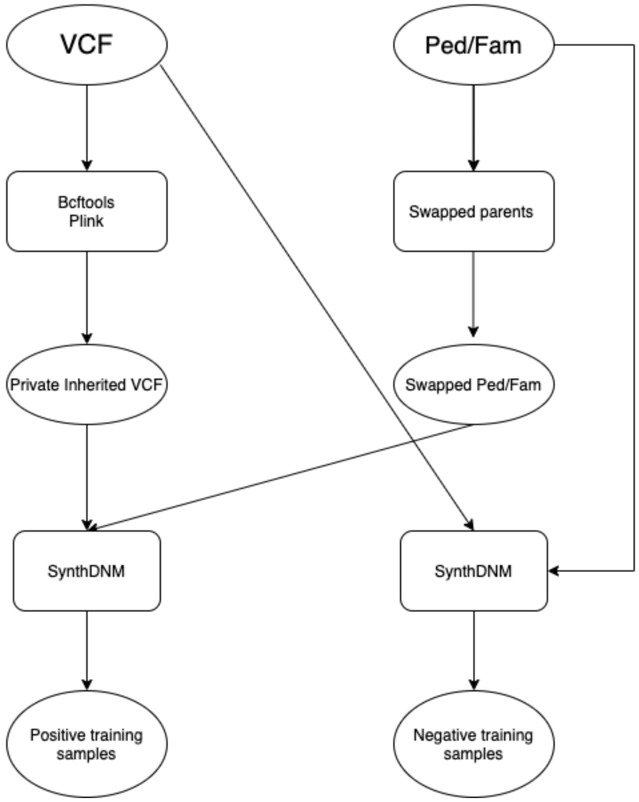


Fig.S2 Workflow of creating training set. For generating positive training samples, input VCF will annotated and run TDT test by Bcftools and PLINK, then obtain private inherited VCF, Ped or Fam file will swapped to swapped-ped/fam file. Using these two files, SynthDNM can generate positive training samples; For generating negative training samples, SynthDNM can using input VCF and Ped/Fam to extracting all putative *de novo* mutations as negative training samples.

**Variants on sex chromosome**

In order to predicting dnms on variants of sex chromosome, we build different training set for different type of variants. For variants on female X chromosome and on male PAR regions of sex chromosomes, we could use the classifier trained on autosome; For variants on male Non-PAR regions, first select the private inherited variants as follows:

| | child | mother | father |
|---|---|---|---|
| chrX | 1\|1 | 0\|1 | **** |
| chrY | 1\|1 | **** | 1\|1 |

Then do the swap and the resulting de novo genotypes should be:

| | child | mother | father |
|---|---|---|---|
| chrX | 1\|1 | 0\|0 | **** |
| chrY | 1\|1 | **** | 0\|0 |

After swapped parents, all the "synthetic DNMs" are label as positive training set, also, all the putative dnms on sex chromosome are label as negative training set as before.

**Training Features**

In our case, for classifier trained on GATK VCFs, all the predictive features, all derived from metrics contained in the VCF, are described below.

parent_ar_max: the maximum of the parental ratios of mutant (child) allele reads to parental allele reads

parent_ar_min: the minimum of the parental ratios of mutant (child) allele reads to parental allele reads

offspring_ar: the ratio in the offspring of mutant allele reads to parental allele reads

parent_dp_max: the max allele depth of parent

parent_dp_min: the minimum allele depth of parent

offspring_dp: the allele depth of offspring

parent_dnm_pl_max: the maximum of the parents' Phred likelihoods for the heterozygous genotype (i.e. the DNM genotype, 0/1)

parent_dnm_pl_min: the minimum of the parents' Phred likelihoods for the heterozygous genotype

offspring_dnm_pl: the child's phred likelihood for a heterozygous genotype

parent_inh_pl_max: the maximum of the parents' Phred likelihoods for the homozygous reference genotype (i.e. 0/0)

parent_inh_pl_min: the minimum of the parents' Phred likelihoods for the homozygous reference genotype (i.e. 0/0)

offspring_inh_pl: the child's phred likelihood for a heterozygous genotype for REF

parent_gq_max: maximum of the parental genotype qualities

parent_gq_min: minimum of the parental genotype qualities

offspring_gq: genotype quality for the child

VQSLOD: log odds ratio of being a true variant versus being false under the trained gaussian mixture model

ClippingRankSum:Z-score From Wilcoxon rank sum test of Alt vs. Ref number of hard clipped bases

BaseQRankSum: Z-score from Wilcoxon rank sum test of Alt Vs. Ref base qualities

FS: phred-scaled p-value using Fisher's exact test to detect strand bias

SOR: Symmetric Odds Ratio of 2x2 contingency table to detect strand bias

MQ: mapping quality

MQRankSum: Z-score From Wilcoxon rank sum test of Alt vs. Ref read mapping qualities

QD: variant confidence/quality by depth

ReadPosRankSum: Z-score from Wilcoxon rank sum test of Alt vs. Ref read position bias

For classifier trained on Deep_variant and Wecall VCFs, here are features we used:

offspring_ar: the ratio in the offspring of mutant allele reads to parental allele reads

parent_dp_max: the max allele depth of parent

parent_dp_min: the minimum allele depth of parent

offspring_dp: the allele depth of offspring

parent_gq_max: maximum of the parental genotype qualities

parent_gq_min: minimum of the parental genotype qualities

offspring_gq: genotype quality for the child

AQ: Allele Quality score reflecting evidence for each alternate allele (Phred scale)

offspring_reference_pl: the child's Phred likelihood for a homozygous reference genotype

offspring_alternate_pl: the child's Phred likelihood for a homozygous alternate genotype

**SynthDNM Classifier Parameter Selection**

As previously mentioned, we use a random forest classifier from scikit-learn, which is a meta estimator that fits a number of decision tree classifiers on various sub-samples of the dataset and uses averaging to improve the predictive accuracy and control over-fitting. Random forest classifiers are governed by n_estimators、max_depth、min_samples_split、min_samples_leaf、min_weight_fraction_leaf、max_features、max_leaf_nodes. Parameter sweeps of each values were performed with balanced class weights and parameters were chosen by optimizing classfication accuracy with cross validation of the training set from gridsearch.

#### Feature importance

Random Forest classifiers include an internal measure of feature importance, which indicates a feature's contribution to predictive accuracy. This is done by randomly permuting each feature and then assessing the resulting degradation of predictive performance for each class, as well as across all classes. Here, we show the feature importance for each class in Figure S3.

Fig.S3

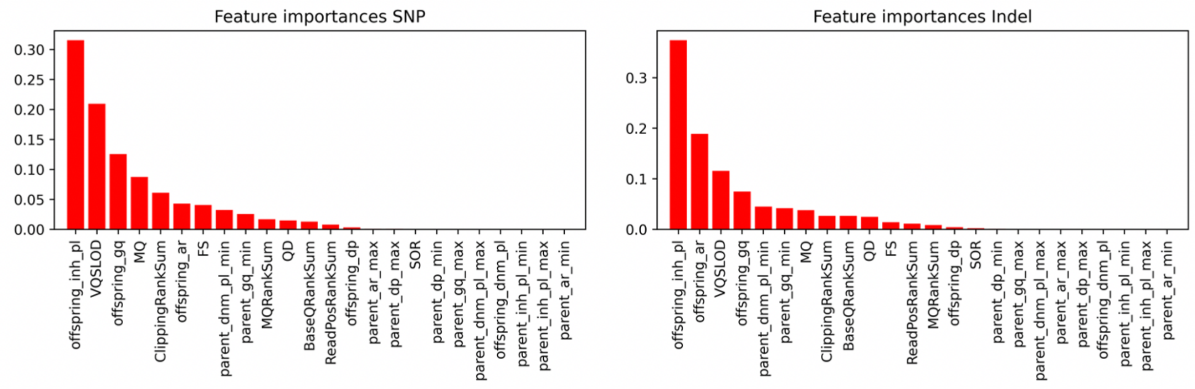


Fig.S3 Feature importance of pre-trained classifier. In our case, For the SNPs, the offspring_inh_pl and VQSLOD, as well as offspring_gq are the most important features for the training set. For the Indels, offspring_inh_pl、offspring_ar and VQSLOD are the most important features for the training set.

#### References


Brandler,W.M. *et al.* (2018) Paternally inherited cis-regulatory structural variants are associated with autism. *Science*, **360**, 327–331.

Gardner,E.J. *et al.* (2019) Contribution of retrotransposition to developmental disorders. *Nat. Commun.*, **10**, 4630.

Koboldt,D.C. *et al.* (2012) VarScan 2: somatic mutation and copy number alteration discovery in cancer by exome sequencing. *Genome Res.*, **22**, 568–576.

Koboldt,D.C. *et al.* (2009) VarScan: variant detection in massively parallel sequencing of individual and pooled samples. *Bioinforma. Oxf. Engl.*, **25**, 2283–2285.

Li,B. *et al.* (2012) A Likelihood-Based Framework for Variant Calling and De Novo Mutation Detection in Families. *PLoS Genet.*, **8**, e1002944.

Michaelson,J.J. *et al.* (2012) Whole-genome sequencing in autism identifies hot spots for de novo germline mutation. *Cell*, **151**, 1431–1442.

Mohanty,A.K. *et al.* (2019) novoCaller: a Bayesian network approach for de novo variant calling from pedigree and population sequence data. *Bioinforma. Oxf. Engl.*, **35**, 1174–1180.

Purcell,S. *et al.* (2007) PLINK: A Tool Set for Whole-Genome Association and Population-Based Linkage Analyses. *Am. J. Hum. Genet.*, **81**, 559–575.

Quinlan,A.R. and Hall,I.M. (2010) BEDTools: a flexible suite of utilities for comparing genomic features. *Bioinformatics*, **26**, 841–842.

Ramu,A. *et al.* (2013) DeNovoGear: de novo indel and point mutation discovery and phasing. *Nat. Methods*, **10**, 985–987.

Wei,Q. *et al.* (2015) A Bayesian framework for de novo mutation calling in parents-offspring trios. *Bioinformatics*, **31**, 1375–1381.

Werling,D.M. *et al.* (2018) An analytical framework for whole genome sequence association studies and its implications for autism spectrum disorder. *Nat. Genet.*, **50**, 727–736.
